# Female investment in terminal reproduction or somatic maintenance depends on infection dose

**DOI:** 10.1101/2022.07.25.501224

**Authors:** Nora K. E. Schulz, Carly M. Stewart, Ann T. Tate

## Abstract

Immune responses are energetically costly to produce and often trade off with investment in reproduction and offspring provisioning. We predicted that hosts can interpret the magnitude of the infection threat from cues like initial bacterial load to adjust reproductive investment into offspring quantity and quality. To test this prediction, we exposed female flour beetles (*Tribolium castaneum*) to naïve or sterile media controls or one of three increasing doses of heat-killed *Bacillus thuringiensis* (Bt). To estimate offspring quantity, we measured the number of eggs and hatched larvae from each maternal treatment group. To estimate offspring quality, we measured egg protein content, development to pupation, pupal weight, and offspring survival against Bt infection. Compared to naïve controls, low and intermediate bacterial doses resulted in lower female fecundity, suggesting a shift to a somatic maintenance strategy. Meanwhile, in the highest-dose group, fecundity was negatively correlated with egg protein content, suggesting a trade-off between offspring quality and quantity that could mask terminal investment based on metrics of quantity alone. Our results underscore the need to account for the magnitude of environmental cue when quantifying plasticity and trade-offs among life history traits, and provide new insight into the trans-generational effects of immune responses.

## Introduction

Parasites and pathogens steal energetic resources from their hosts and damage host tissues. The energy that hosts allocate to resist infections, repair damage, and maintain homeostasis can come at the expense of other life history traits like reproduction. Sometimes, it can be in the interest of host fitness to reproduce quickly and risk subsequent disease-induced mortality rather than invest in costly immune defence mechanisms [1–3]. The terminal investment hypothesis predicts that if the likelihood of imminent death is relatively high, individuals can opt to maximize current reproductive output (possibly at the expense of future reproduction) and thus maximize their fitness [1,4]. For example, snails that have been exposed to a castrating parasite partially compensate for their expected loss of fecundity by increasing egg laying rates [5]. Alternatively, if the danger to future reproduction is less severe, individuals may invest in somatic maintenance, as when mealworm beetles temporarily restrained their reproduction after a relatively avirulent bacterial immune challenge [6] or as when female dung beetles reduced their reproductive output after immune activation [7]. By reducing offspring production in favor of survival and immunity over the short term, individuals gain a longer potential reproductive period.

Mounting evidence suggests that reproductive strategies are not fixed and intransigent, such that a genotype or species will always resort to either terminal investment or somatic maintenance upon infection. Among flour beetle populations, for example, the induction of an immune response led to either a survival benefit for the mothers or an increase in reproduction after secondary pathogen exposure [8]. More generally, reproductive effort appears to respond dynamically to factors that affect residual reproductive potential, including host age and the availability of energetic resources [9]. For example, both infection and old age can cause a shift to terminal investment in burying beetles [10,11]. In blue footed boobies, younger males reduce their reproductive output in response to an immune challenge, while old males almost double theirs [12], suggesting age-related plasticity. Fruit flies deprived of protein mainly exhibit variance in overall survival, suggesting that they allocate limited protein to staying alive, while protein supplementation spurs the flies to shift investment to maximizing fecundity [13]. Thus, reproductive strategy is likely plastic and sensitive to the life history and environmental context of the host.

Parasite virulence and initial infection dose also influence the probability of host survival, the length of the reproductive lifespan, and – to the extent that the host can accurately interpret information about the infection – the relative costs and benefits of investing time and energy into immunity or reproduction [9,14]. For example, increasing doses of the bacterium *Pseudomonas syringae* was associated with increased per-capita reproductive rates in pea aphids [15], although reproductive investment outcomes in aphids also depend on the type of challenge (e.g., bacterial strain) and aphid genotype [14]. Young male crickets reduced mating effort regardless of infection dose, while older males increased mating effort at the highest doses [16]. Taken together, these studies suggest that host physiology likely interacts with cues about the potential for parasite-induced pathology to modulate relative investment in somatic maintenance and fecundity.

When weighing strategies that maximize individual fitness, however, it is important to remember that the number of offspring is only part of the equation – those offspring must also be able to survive to reproduce. If parents and offspring share the same environment, the new generation is likely to encounter the same parasites, and their chances of survival would likely benefit from additional provisioning from the parent. While the somatic maintenance versus terminal investment dilemma focuses on the dimension of offspring quantity, investment in offspring quality likely co-varies with strategy. While parental care is limited in many insects, parents may improve the odds of survival for their offspring by providing energetic resources to the eggs or accelerating offspring development to reduce the time spent in vulnerable stages [17,18].

Parents may also invest in offspring quality through trans-generational immune priming, as observed in a variety of insect species (as reviewed in [19–22]). In the mealworm beetle *Tenebrio molitor*, for example, mothers exposed to bacteria are capable of bequeathing antimicrobial peptides to their eggs [23], and the proportion of protected eggs was positively associated with fecundity – up to a point. As fecundity continued to increase, however, a clear trade-off between offspring quantity and provisioning was observed [23]. Just as with reproductive investment, priming investment may depend on the strength of the parental cue (i.e., dose). In African armyworms primed with a virus, low parental doses resulted in increased offspring survival of infection, while high doses had the opposite effect, demonstrating the dose dependency of trans-generational immune priming [24]. Although the mechanisms behind trans-generational priming remain debated, the additional investment into immune responses that it requires should be costly for both mother and offspring and therefore incompatible with terminal investment. Underscoring this intuition, a stochastic agent-based model of the evolution of somatic maintenance and terminal investment predicted that the presence of trans-generational priming in a system should strongly favour somatic maintenance, even under conditions when terminal investment would otherwise be favoured [25].

Taken together, these observations suggest that both fecundity and investment in offspring quality respond plastically to the magnitude of immune challenge (the dynamic terminal investment threshold hypothesis, [9]) and that investments in offspring quality and quantity likely trade off. Therefore, we predicted that insects exposed to lower doses of bacteria would shift to somatic maintenance and produce offspring that were fewer in number but better provisioned, while higher doses would stimulate terminal investment and low-quality offspring.

To test these predictions, we employed flour beetles (*Tribolium castaneum*), a stored product pest beetle and model organism frequently used in studies of trans-generational immune priming [26–28]. We challenged them with increasing heat-killed doses of the natural entomopathogen *Bacillus thuringiensis* (Bt) to change the strength of the danger signal while controlling for microbial pathology. To estimate maternal fecundity, we measured both the number of eggs laid and the number of surviving larvae for several weeks after maternal immune challenge. To measure offspring quality in response to each maternal treatment, we measured egg protein content, offspring development time, pupal mass, and survival rates following live bacterial infection. Our results suggests that while reproductive responses to infection are dose dependent, dose only marginally affects offspring quality. These results reveal the extent to which offspring quality and quantity covary across environments and provide support for using dose-response designs in experiments meant to quantify costs and trade-offs associated with immune investment and trans-generational effects.

## Materials and Methods

### The model system

In this study we used a lab stock population of red flour beetles (*Tribolium castaneum*) that were collected from a grain elevator in Pennsylvania, USA in 2013[29]. Beetles were kept in non-overlapping generations on sterilized whole wheat flour supplemented with 5% brewer’s yeast in the dark at 30°C and 70% humidity.

For the immune challenge we used a strain of *B. thuringiensis berliner* (Bt, ATCC 55177) featured in a previous dose response infection experiment [30]. Bt is a natural pathogen of beetles, it elicits immune responses even when heat-killed, and it is capable of stimulating trans-generational priming in the beetles [26–28,31]. Bt is an obligate killer that naturally infects via the oral route, but also produces reliable infection outcome and priming results when introduced septically directly into the haemolymph [28,31,32].

### Production of the maternal generation

To produce the experimental parental generation, we set up five replicates of ~100 beetles each from our stocks. Each replicate consisted of a Petri dish (100mm diameter) half filled with the standard diet. Beetles laid eggs for three days and then were removed from the dish. Once most of the offspring had pupated, we determined their sex, and individualized them into flat bottom 96-well plates. We supplied individuals that eclosed with a small amount of diet added to the well. When all individuals had reached sexual maturity, i.e., four days after the last individuals had eclosed, we performed the maternal exposure treatment.

### Maternal bacterial exposure

For the maternal bacterial exposure treatment, we grew an overnight Bt culture in Luria-Bertani broth from a glycerol stock stored at −80°C. We washed the cultures twice with insect saline and adjusted the OD600 to 1.0 by resuspending bacteria in insect saline. In addition to naïve (handled but not injected) and sterile saline-injected controls, we made two tenfold dilutions to create three priming doses: high dose (undiluted bacteria suspension with ~3*10^8^ colony forming units (CFU/mL), intermediate dose (1:10 dilution), and low dose (1:100 dilution). Afterwards we heat-killed the bacteria by exposing them to 90°C for 30 min. Samples of live and heat-killed bacteria spread onto LB agar plates and incubated overnight provided us with the number of CFUs/mL in our priming doses and confirmed that the heat had killed all cells, respectively. We randomly assigned all sexually mature females to one of the five priming treatment groups.

We carried out the priming as previously described, but without anaesthetizing the beetles [26]. In short, we pricked beetles with an ultrafine needle between occiput and pronotum after dipping it first in sterile insect saline and then into heat-killed bacteria solution according to treatment. Beetles of the naïve group were handled in the same manner without being pricked with the needle. Afterwards, we put each female into an empty well of a new 96-well plate and recorded their survival after 24h.

### Mating and egg laying

For the mating, we put each of the surviving treated females together with one untreated male into a small Petri dish (50mm diameter) half filled with standard diet (n=30). After 48h, all pairs were collected and moved to new petri dishes that contained the standard diet of flour and yeast, which we pre-sieved to enable egg collection. After a 24h oviposition period, we transferred all pairs to individual dishes containing standard diet to avoid any reduction in egg production due to malnourishment from the pre-sieved flour, which contains less protein because the sieving removes a large proportion of the yeast. In total, we collected eggs six times in this manner over a period of 15 days.

### Fecundity measurements

For all six oviposition periods, we counted the number of eggs produced for each individual mating pair (including all females and excluding females that died during oviposition periods). For oviposition periods 1, 3 and 5 we additionally counted the number of live larvae two weeks post oviposition. All pairs in which the male died, an individual escaped, or which did not produce any offspring in any of the egg lays were excluded from analysis (excluded pairs out of 30 per treatment: naïve= 4, wounding= 5, low= 0, intermediate= 3, high= 6).

### Egg protein content

We measured total protein content in a subsample of eggs to approximate maternal energetic investment into offspring. For this we used eggs produced during the second oviposition period from females that had at least laid ten eggs (minimum number needed for reliable results, n=16 pools of 10-36 eggs per treatment). We assume that higher protein content signals better quality offspring, which has more resources available for growth and development [33]. We adapted a Bradford assay originally used in the analysis of *Drosophila melanogaster* eggs [34]. In brief, eggs were stored at −20°C until the day of the assay, whereupon we sieved the eggs again, removed all larger diet particles, recounted them, and transferred them to clean 1.5ml tubes. We washed the eggs twice with phosphate buffered saline (PBS). Then, we added 100μl PBS and two tungsten beads (2.4mm diameter) to each tube and homogenized the eggs in a tissue lyser with 40 oscillations per second for three minutes. In a new tube, we added 30μl of the egg lysate to 500μl of Coomassie blue dye. We measured absorbance at 595nm for two 200μl technical duplicates for each sample. A standard curve on twofold dilutions of albumin standard ranging from 2mg/ml to 0.125mg/ml was used to calculate the total protein content within each sample. We divided the measured protein content by the number of eggs in the sample to obtain measurements of average protein per egg.

### Offspring development and pupal mass

We followed the larval offspring from oviposition 1, 3 and 5 through their development and recorded the number of individuals that had pupated at 16 days after oviposition, a time point chosen to maximize statistical power from cross-sectional data as on average ~50% of individuals from this population will have pupated by then. At day 16, there were some pupae in all groups, and none had reached 100% pupation.

On day 16 and 17 post oviposition, we recorded the weights of one randomly picked freshly formed pupa per day and parental pair. We used the mean weight of the two pupae from the same parents in the following analysis. This was done for the three oviposition periods (1, 3 and 5).

### Offspring survival of infection

To test for potential effects of transgenerational immune priming, we septically infected offspring from egg lays 1, 3 and 5 with an LD30 dose (~3*10^8^ CFU/mL) of Bt and recorded their survival 24h later. We collected three larvae from each parental pair that had produced at least five live offspring (which excluded 2-5 pairs per oviposition period and treatment) for the Bt infection treatment. Additionally, we collected twelve larvae from random pairs (1 larvae/pair) for the wounding control treatment. For the immune challenge, we grew an overnight culture of Bt as described above. From this culture we started new cultures in 3mL LB, which we harvested after two hours when they still were in the logistic growth phase. The OD at 450nm for both log phase and overnight cultures was adjusted to 1 and both cultures were combined in a 1:1 ratio, which leads to the most consistent mortality rates. We then centrifuged and resuspended the bacterial suspension in 200 μl insect saline. We plated an aliquot of the insect saline and serial dilutions of the bacteria suspension on LB agar plates to confirm sterility of the insect saline and estimate infection dose by counting CFUs after incubation (infection dose ranged between 2.5-3*10^8^ CFUs/mL for the three oviposition periods).

For the septic infection, we used larvae that were 14-15 days post-oviposition. Larvae received a prick with an ultra-thin needle, dipped first in insect saline and then in the respective treatment suspension (either insect saline for wounding control or bacteria suspension for the infection challenge treatment). We pricked larvae laterally between the second and third to last distal segment. After treatment, larvae were placed individually into empty wells of a 96 well plate and kept at 30°C for 24 hours until survival was assessed.

### Statistical analysis

We conducted all statistical analyses in R (version 4.1.0, [35]) and RStudio (version 1.4.1717, [36]) using packages lme4 [37] and MASS [38] to produce GLMs and GLMMs. We investigated the survival of females after priming treatment in a Cox Proportional Hazard analysis, after confirming the assumptions of proportionality were not violated, using the naïve group as the base level.

All egg and larval count data were over-dispersed and followed a negative binomial distribution of error terms. Using a negative binomial GLMM with breeding pair as a random effect to account for repeated measurements, we first tested for an interaction between Bt exposure treatment and oviposition period (eggs ~ treatment * ovp + (1 | pair). Because the interaction term was not significant, we then combined all counts for individual oviposition periods into one value. For the egg counts, we ran analyses on two data sets, one including females that died during the six oviposition periods and one excluding data from families where the female died before the final egg lay. In both analyses, and for the analysis of larval counts, we applied the same GLM with treatment as the sole fixed effect (number of eggs or larvae ~ treatment). Additionally, we tested whether the hatching rate interacted with maternal treatment in a GLM with number of larvae as the dependent variable and treatment and number of eggs as fixed effects (larvae ~ treatment * eggs, with a negative binomial error term).

The egg protein content data followed a Gamma distribution and was analysed with a GLMM with exposure treatment as a fixed effect and block (96 well plate on which the OD measurements were conducted) as a random effect (protein per egg ~ treatment + (1|block)). For each maternal exposure treatment, we tested for a significant linear correlation between per egg protein content (ovp 2) and number of eggs laid (ovp1-6) as well as number of larvae (ovp 1, 3 and 5). We corrected for multiple testing using the Benjamini-Hochberg method with a false discovery rate set to 0.1 [39].

We tested for the effects of maternal exposure treatment on the proportion of pupae at day 16 post oviposition (binomial error distribution) with a GLMM with treatment as fixed effect and oviposition period and parental pair as random effects to account for repeated measurements (proportion of pupae ~ treatment + (1 | ovp) + (1 | pair)). The same random effects were included in the linear mixed model testing the effect of maternal treatment on pupal weight (weight ~ treatment + (1 | ovp) + (1 | pair)). Additionally, we investigated whether there were significant linear correlations between the offspring quality measures (pupation and pupal weight) and offspring quantity (eggs laid (ovp1-6) and live larvae (ovp 1, 3 and 5).

Finally, to investigate offspring survival of septic infection and test for the presence of trans-generational priming, we applied a GLM with a binomial error distribution and maternal exposure treatment and oviposition period as main effects and their interaction (proportion alive ~ treatment * ovp).

## Results

### Female fecundity after Bt exposure treatment

To estimate how female fecundity is affected by exposure to different doses of heat-killed bacteria, we recorded the survival of females over the six oviposition periods and counted their laid eggs and larval offspring. Overall, less than 7% of females died from the treatment itself in the first 24 h post treatment. These deaths were evenly distributed across the five treatment groups (1-3 individuals per treatment) and can most likely be attributed to trauma during wounding and handling. Despite not being exposed to live bacteria, more than 20% of females from the three treatment groups exposed to heat-killed bacteria died over the 18-day mating and oviposition period (Figure 1). According to a Cox-proportional hazard analysis (Suppl. Table 1), the groups receiving a low and intermediate dose of heat-killed bacteria had significantly higher mortalities than the naïve control (low dose: HR=8.21, p=0.047; intermediate dose HR=8.5, p=0.045), while the high dose treatment and wounding control group did not differ significantly from the naïve group (high dose: HR=6.84, p=0.075; wounding control: HR=2.11, p=0.54). The high dose mortality was very similar to the mortality of the other two heat-killed doses, however, suggesting that the lack of significance may represent a slightly underpowered sample size rather than the lack of a biological effect (Figure 1, Suppl. Table 1).

**Figure 1.**
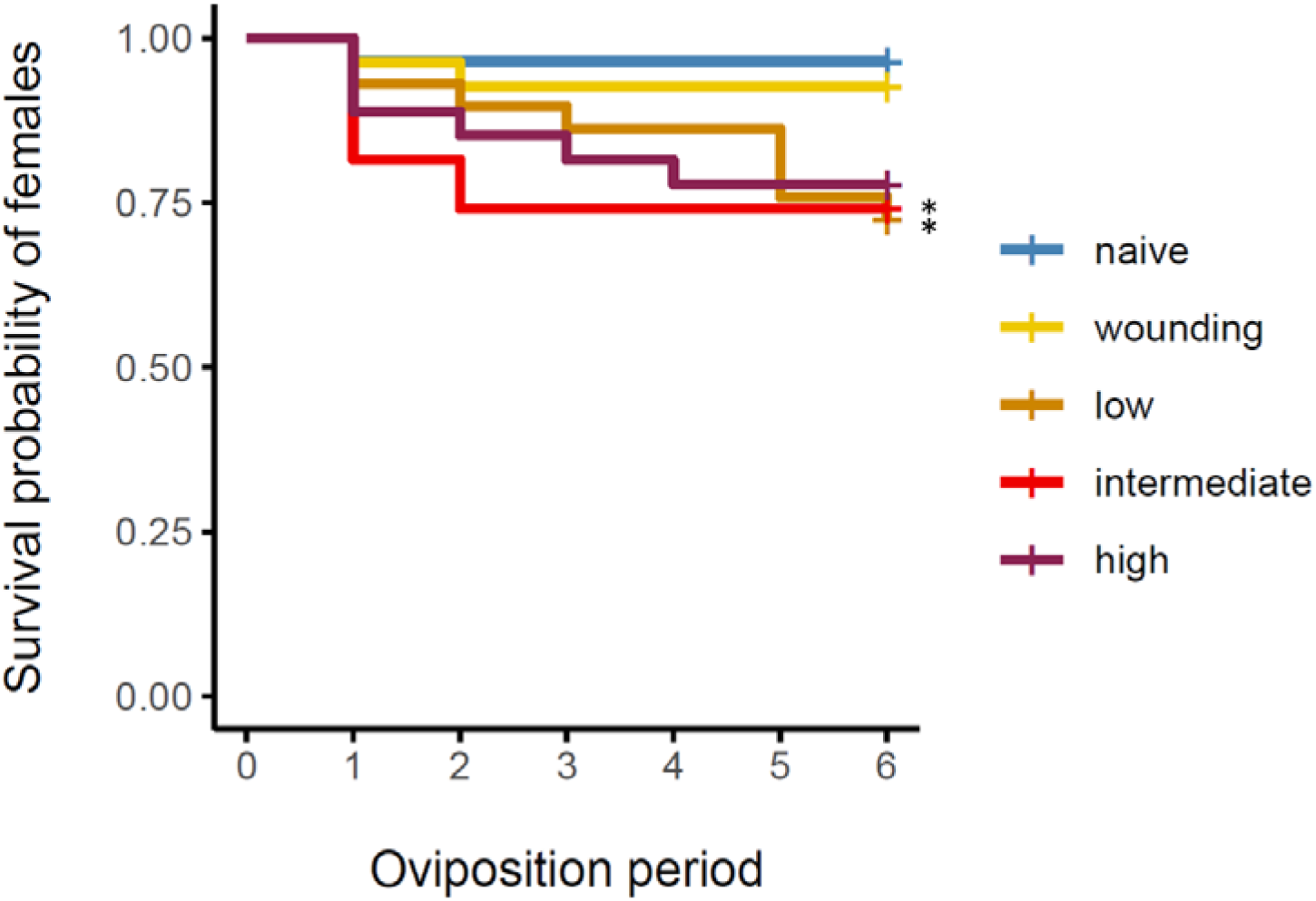
Survival probability for the mated females according to the five treatment groups (naïve n=28, wounding control n=27, low Bt dose n=29, intermediate Bt dose n=27, high Bt dose, n=27). Survival was recorded over six oviposition periods, each three days apart. Asterisks indicate significant difference from naïve treatment group with p<0.05.

### Fecundity after Bt exposure

We counted eggs produced by each mating pair for each of the six oviposition periods. We combined all counts for each pair into one value because the egg-laying rate did not vary significantly over time between the treatment groups (Suppl. Figure 1, Suppl. Table 2). Egg production was significantly reduced in the group that had received the low Bt dose compared to the naïve control (Estimate=-0.26, p=0.04, Suppl. Figure 2, Suppl. Table 2). To exclude the possibility that the reduced egg laying rate from females that died during the experiment was the sole contributor to this finding, we ran the same analysis on the data set containing only egg counts from pairs where the female had survived past the sixth oviposition period. After excluding these females (Figure 1), there was a significant reduction in egg laying rate in low and intermediate dose treatment groups (low dose: Estimate=-0.185, p=0.016; intermediate dose: Estimate=-0.242, p=0.002, Figure 2A, Suppl. Table 2). Females that received a low or intermediate dose of heat-killed bacteria produced on average 22% and 23% fewer eggs than the naïve group. Egg numbers for the wounding control and high dose groups were only marginally reduced by 7% and 6% respectively and did not differ significantly from those laid by the naïve group (wounding control: Estimate=-0.108, p=0.143; high dose: Estimate=-0.047, p=0.55, Figure 2A, Suppl. Table 2).

**Figure 2.**
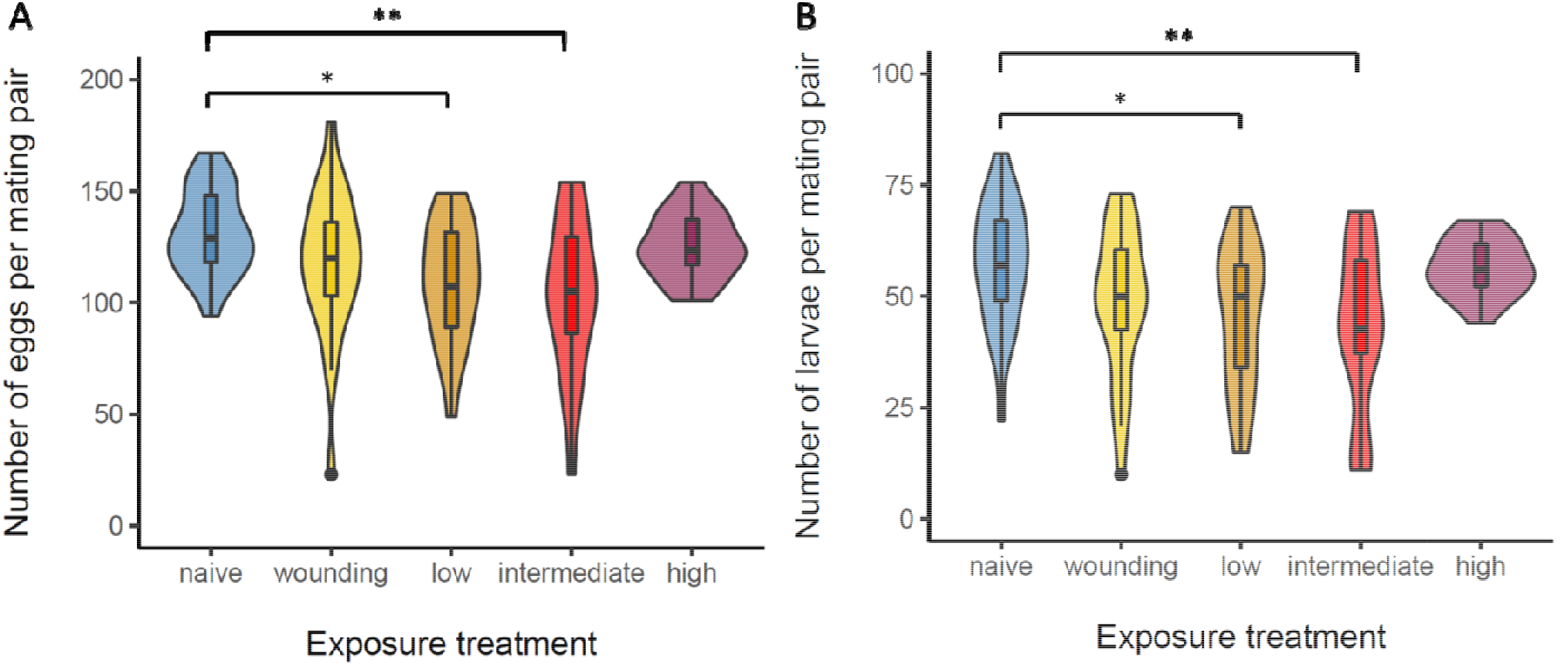
Fecundity after maternal sham infection (for females that survived until the end of the experiment and produced offspring across all oviposition periods). **A)** total number of eggs produced by individual pairs over six 24h oviposition periods (naïve n=25, wounding control n=23, low dose of heat killed Bt n=20, intermediate dose n=19, high dose n=18). **B)** Hatched and surviving larvae per mating pair. Data from oviposition period 1, 3 and 5 combined (naïve n=25, wounding control n=23, low dose of heat killed Bt n=21, intermediate dose n=20, high dose n=19). Asterisks indicate significant differences (*=p<.05, **=p<.01).

For oviposition periods 1, 3 and 5, we let the eggs develop and counted live larvae produced by each female two weeks later. We also combined these counts into one total value per female for the analysis. Similar to the egg count, females that received a low or intermediate dose had significantly fewer larval offspring than the naïve group (low dose: 24% reduction, Estimate=-0.23, p=0.018; intermediate dose: 25% reduction, Estimate=-0.288, p=0.005), while there was no significant difference between the naïve group and the wounding control (15% fewer larvae, Estimate=-0.162, p=0.094) or the group that received the high dose of heat killed bacteria (7% fewer larvae, Estimate=-0.017, p=0.871) (Figure 2B, Suppl. Table 2). None of the four treatment groups differed in their hatching rate (number of larvae/number of eggs) from the naïve group (Suppl. Figure 3, Suppl. Table 2).

### Offspring quality: egg protein content, pupation rate and pupal weight

We evaluated the total protein content per egg for eggs laid during the second oviposition period as an indirect measure of offspring quality. Neither the wounding nor any of the three infection treatments significantly altered average offspring egg protein content compared to the naïve control (Figure 3A, Suppl. Table 3). However, we did see evidence of interaction effects between egg number and egg protein content. While egg number did not significantly predict protein content in eggs from naïve (R^2^=0.09, p=0.14) or saline-exposed mothers (R^2^=0.03, p=0.24), those from a mother receiving a low bacterial dose exhibited a positive correlation between the two metrics (R^2^=0.29, p=0.019), potentially revealing a quality gradient of reproductive potential among the females (Figure 3B). In the high dose group, on the other hand, there was a significant negative relationship between egg quantity and protein content (R^2^=0.23, p=0.036), suggesting a forced trade-off in reproductive investment (Figure 3B).

**Figure 3.**
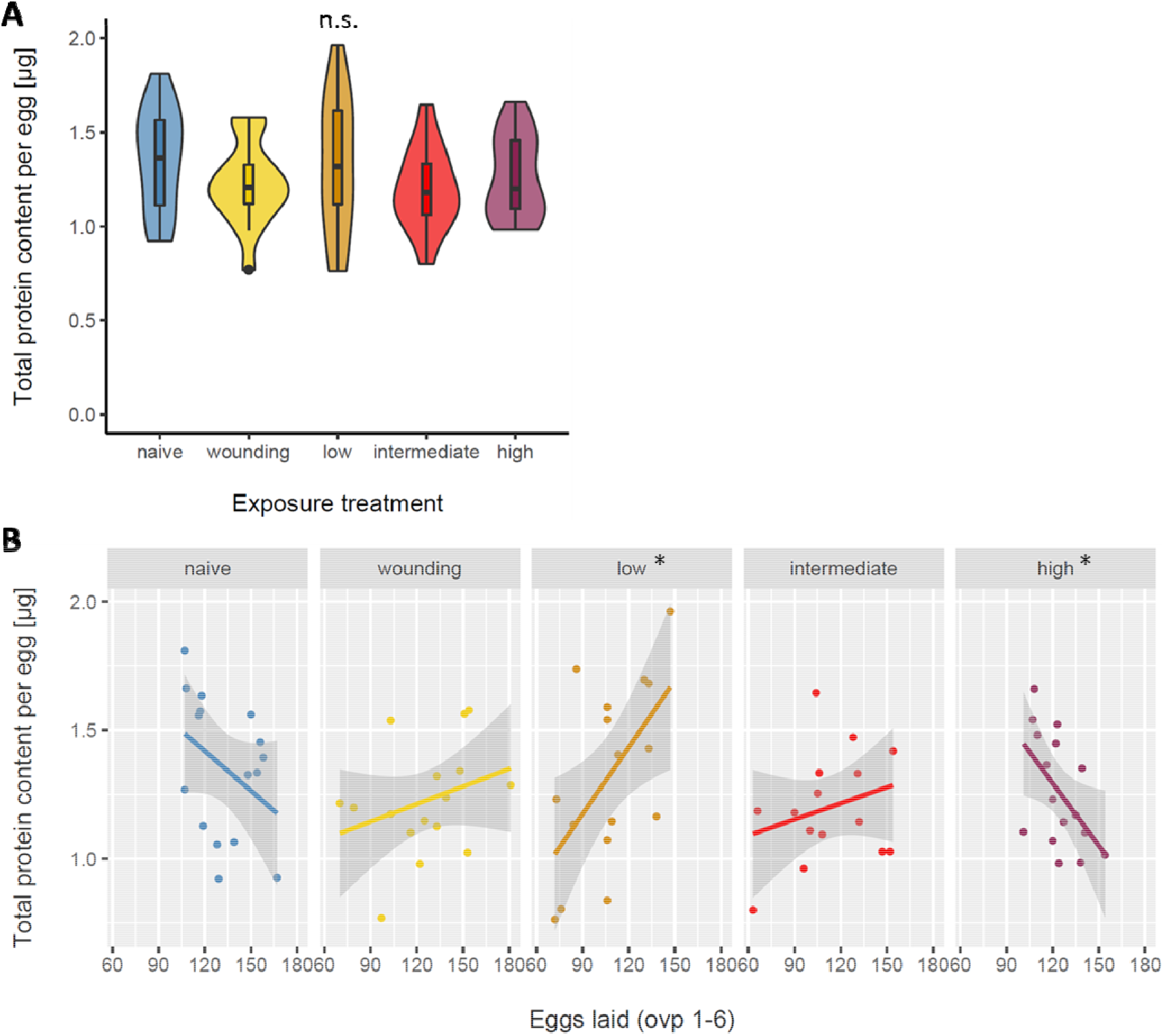
Egg quality after maternal Bt exposure **A)** Total protein content per egg [μg] for each maternal exposure treatment (naïve control n=16, wounding control n=16, low dose of heat killed Bt n=15, intermediate dose n=16, high dose n=16). Bradford protein assay was run on 10-36 pooled eggs per breeding pair. **B)** Correlation between egg protein content per egg (ovp 2) and eggs laid (ovp 1-6) for each of the five maternal exposure treatments (n= 15-16 mating pairs). Asterisks indicate significant correlations with p<.05 (Suppl. Table 4)

Offspring developmental speed and size are reliable indicators of their quality [17,18,40]. Earlier pupation means the individual spends less time in its most vulnerable state, and larger size is associated with higher reproductive fitness. We measured developmental speed by recording the proportion of individuals that had pupated sixteen days post oviposition. There was a significantly higher proportion of pupae in the wounding control (Estimate=-0.466, p=0.012) and intermediate dose treatment (Estimate=-0.793, p<0.001) than in the naïve group (Figure 4A, Suppl. Table 5). Low (Estimate=-0.319, p=0.093) and high dose groups (Estimate=-0.238, p=0.216) were not significantly different from the naïve control (Figure 4A, Suppl. Figure 4, Suppl. Table 5).

**Figure 4.**
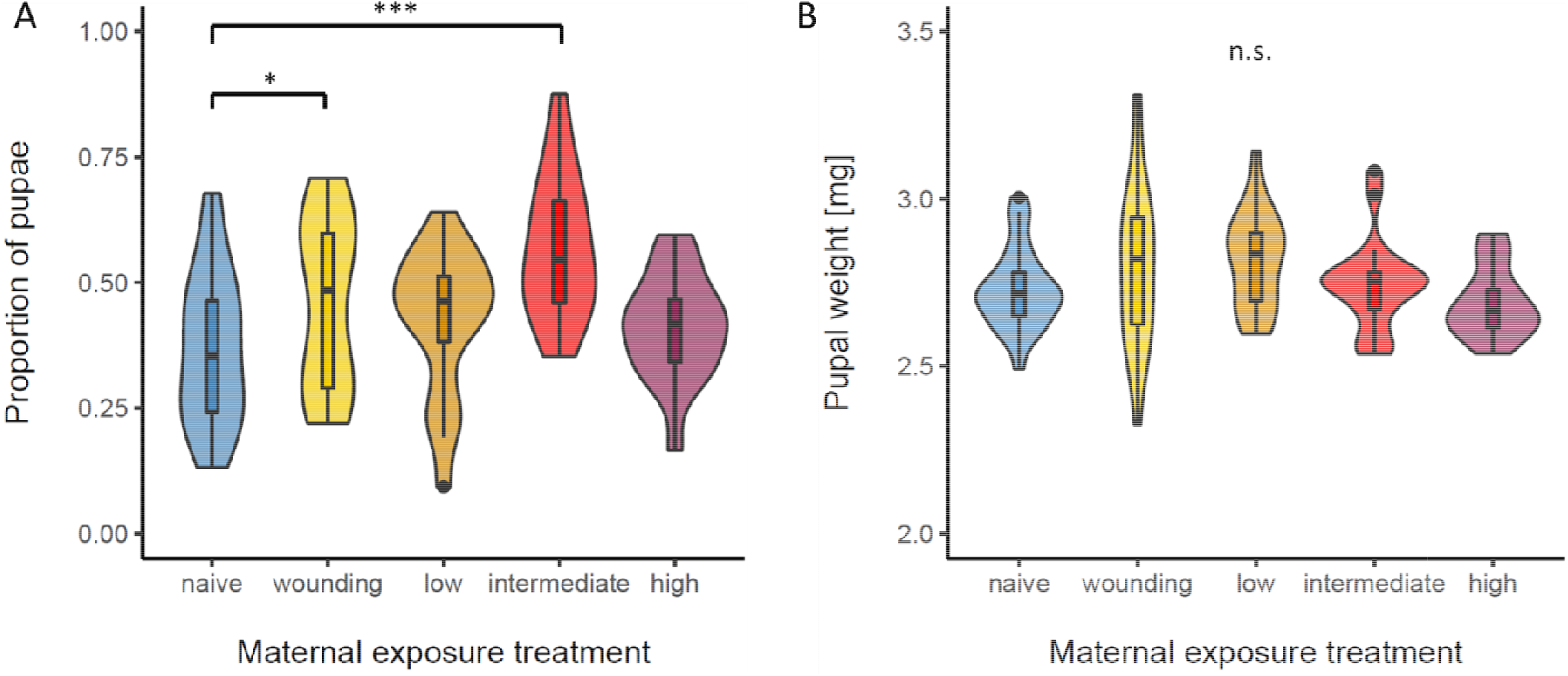
Offspring quality after maternal Bt exposure **A)** Pupation rate (Proportion of pupae to total live offspring) on day 16 post oviposition for the five maternal treatment groups (naïve n=25, wounding control n=22, l low dose of heat killed Bt n=20, intermediate dose n=18, high dose n=18). Means for each breeding pair combined from oviposition period 1, 3 and 5. **B)** Pupal weights [mg] for the five maternal treatment groups (naïve n=25, wounding control n=23, ow dose of heat killed Bt n=20, intermediate dose n=20, high dose n=18). Mean weights calculated for two pupae per pair and oviposition period 1, 3 and 5. Asterisks indicate significant differences with *=p<.05 and ***=p<.001; n.s.=not significant.

For each mating pair, we randomly chose two offspring pupae and weighed them. Pupae did not significantly differ in their weight for any of the four treatments when compared to the naïve group (Figure 4B, Suppl. Figure 4, Suppl. Table 5).

### Offspring survival of septic infection

We septically infected offspring larvae from oviposition period 1, 3 and 5 with a potentially lethal dose of *B. thuringiensis*. All individuals from the sterile saline control injection survived (n=6 per oviposition and maternal treatment). Survival of infection varied between 66%-91% among the oviposition periods (Suppl. Table 6). However, there were no significant differences in survival between any of the four maternal treatment groups and the naïve control (Suppl. Table 7).

## Discussion

Theory predicts that host resource allocation to physiological processes like survival and reproduction should evolve to be sensitive to the magnitude of threats in the environment [14,25,41]. In the case of allocation trade-offs between immunity and reproduction, we predicted young female beetles with their whole reproductive period ahead of them would prioritize survival upon perceiving a surmountable threat like a mild infection, while a more existential threat cued by a high bacterial dose would promote terminal investment in reproduction. Our results suggest that female responses to the inactivated pathogens were indeed dose dependent. Females exposed to a low or intermediate dose of heat-killed bacteria produced fewer offspring, suggesting the adoption of a somatic maintenance strategy that prioritized survival and immune investment over reproduction. Females exposed to the highest bacterial dose did not exhibit the same reduction in fecundity but did reveal a trade-off between offspring quantity and quality, as estimated by the average protein content of eggs. This suggests an effort to prioritize fecundity despite low resources, partially consistent with a terminal investment strategy. To the best of our knowledge, this study is the first to investigate the impact of exposure dose on the sensitivity of maternal investment into offspring quantity and quality, and sheds new light on the evolutionary drivers of plasticity in resource allocation to survival and reproduction.

Previous work investigating reproductive plasticity in response to infection dose have suggested that gross metrics of offspring quantity after exposure are correlated to the residual reproductive period of the parent [15,16,42]. In a study of pea aphids infected with an ultimately lethal bacterium, for example, the lowest initial dose spurred a decrease in reproductive output. Aphid fecundity recovered as dose subsequently increased, suggesting that a shift to somatic maintenance at low doses was replaced by terminal investment as infection-induced mortality became more certain [15]. Aphid reproductive investment never became significantly greater than the fecundity of controls, however, suggesting that microbe-induced pathology could be inducing side-effects like damage to reproductive tissues that obscure the true magnitude of terminal investment. Even though we avoided substantial microbe-induced pathology by using heat-killed bacteria, the fecundity of our beetles demonstrated a similar U-shaped trajectory with increasing doses. Therefore, our results suggest that the magnitude of the cue alone is sufficient to warn of variation in survival probability without continuous feedback from subsequent infection dynamics within the host, potentially through the threshold-like activation of a generalized stress response as observed in the nematode *C. elegans* [43].

As a proxy for host fitness, fecundity is often the most straightforward trait to measure and interpret. However, fitness is maximized only if those offspring are also viable and competitive in their environment. Therefore, increased investment in offspring quality that comes at the expense of current or later fecundity and survival could also fall under the umbrella of terminal investment because it temporarily boosts the production of viable offspring [44,45]. We measured several potential indicators of offspring quality, including egg protein content, development rate, pupal weight, and survival against infection, all of which are positively associated with quality. Our results suggested that while individuals exposed to sham or low doses might produce larvae that develop faster (potentially to escape a hostile environment) [28], there were no other consistent trends of offspring quality variation across exposure dose. When considering the interaction of fecundity and egg protein content, however, we did discover that the relationship between offspring quality and quantity shifts across dose. Mothers exposed to the low dose, for example, were primarily aligned across a maternal quality gradient, where some mothers could produce lots of well-provisioned eggs while others struggled in both metrics. Alternatively, this could reflect natural variation in reproductive strategy, where some mothers strongly prioritized maintenance while others prioritized reproduction to yield a modest net effect. As the dose increased, however, the interaction effect reversed, such that mothers could produce more eggs or well-provisioned eggs but not both. This likely reflects a high energetic cost of the initial immune response [46,47], leaving mothers with few resources to invest in any other trait even if they were inclined to compensate with reproduction.

Because flour beetles are capable of trans-generational immune priming and we successfully measured priming in this beetle population when we first collected them in 2013 [29], we expected offspring from exposed mothers in this study to exhibit increased survival after live Bt infection relative to those born to unexposed mothers [22,27,48]. Alternately, if exposure forced a decrease in maternal provisioning and offspring quality, we would expect to see offspring from exposed mothers become more susceptible to infection [49]. However, we observed no consistent phenotype that would indicate either a priming response or a more general shift in offspring quality. This is not unduly surprising, as the priming phenotype varies across natural populations [27] and its maintenance over evolutionary time may be unstable in natural or laboratory populations [50,51]. It is also possible that any net phenotypes were cancelled out by the opposing forces of priming, which could increase with dose [52], and poor offspring provisioning, which could get more severe at higher doses. Future work would benefit from using a beetle strain with a strong priming phenotype to try to disentangle these potential explanations.

Despite using heat-killed bacteria, which should minimize microbe-induced pathology, we observed increased mortality in female beetles injected with heat-killed bacteria over the eighteen days following the exposure. It is possible that the combination of the stress brought on by repeated mating and the costs of the activated immune responses led to this increased death rate. Mating alone, under monogamous and more so under polyandrous conditions can lead to significant mortality within two weeks in *Tribolium* beetles [53]. Another study found that mated females survived an infection with *B. thuringiensis* at lower rates than virgin beetles [54]. Therefore, our results demonstrate that the combined costs of continuous mating and activated immune responses are of high significance even in the absence of live pathogens. Fortunately, the mortality rates in our study were similar across heat-killed doses, meaning that we do not expect the observed fecundity and provisioning differences among these groups to be attributable to survivorship biases.

### Conclusion

When faced with the threat of a lethal infection, an individual has multiple options to optimize fitness against the twofold costs of reproduction and immune responses. In line with a terminal investment strategy, the infected individual could increase reproduction if death is imminent to maximize fitness. If the risk of death is less severe, the host might invest in somatic maintenance and fight off the infection, potentially reducing reproduction in the meantime. It follows that the severity of infection should affect the outcome of this trade off, meaning higher infection doses should lead to an increase in reproduction while with lower doses we should be able to observe an increased investment in immunity at the expense of reproduction. Our results broadly align with these predictions, but also emphasize the importance of measuring both offspring quality and quantity as metrics of reproductive investment since these can trade off and obscure the true effect. Finally, these results add to the growing evidence that infection dose can qualitatively impact the observation of life history traits and trade-offs [15,16,52], and underscore the importance of dose-response designs in future infection studies.

## Authors’ contributions

AT conceived the study. NS and AT designed the experiment. NS and CS collected the data; NS analysed the data. NS and AT wrote the manuscript. All authors contributed critically to the drafts and gave final approval for publication.

## Data availability statement

All data is included in the supplemental materials.

## Acknowledgements

We like to thank Siqin Liu for her experimental help and Faith Rovenolt and Destane Garrett for pilot work that informed the experimental design. This work was supported in part by Alfred P. Sloan Foundation award #FG-2020-12949 to A.T.T.

