## Supplemental_materials for "Female investment in terminal reproduction or somatic maintenance depends on infection dose"

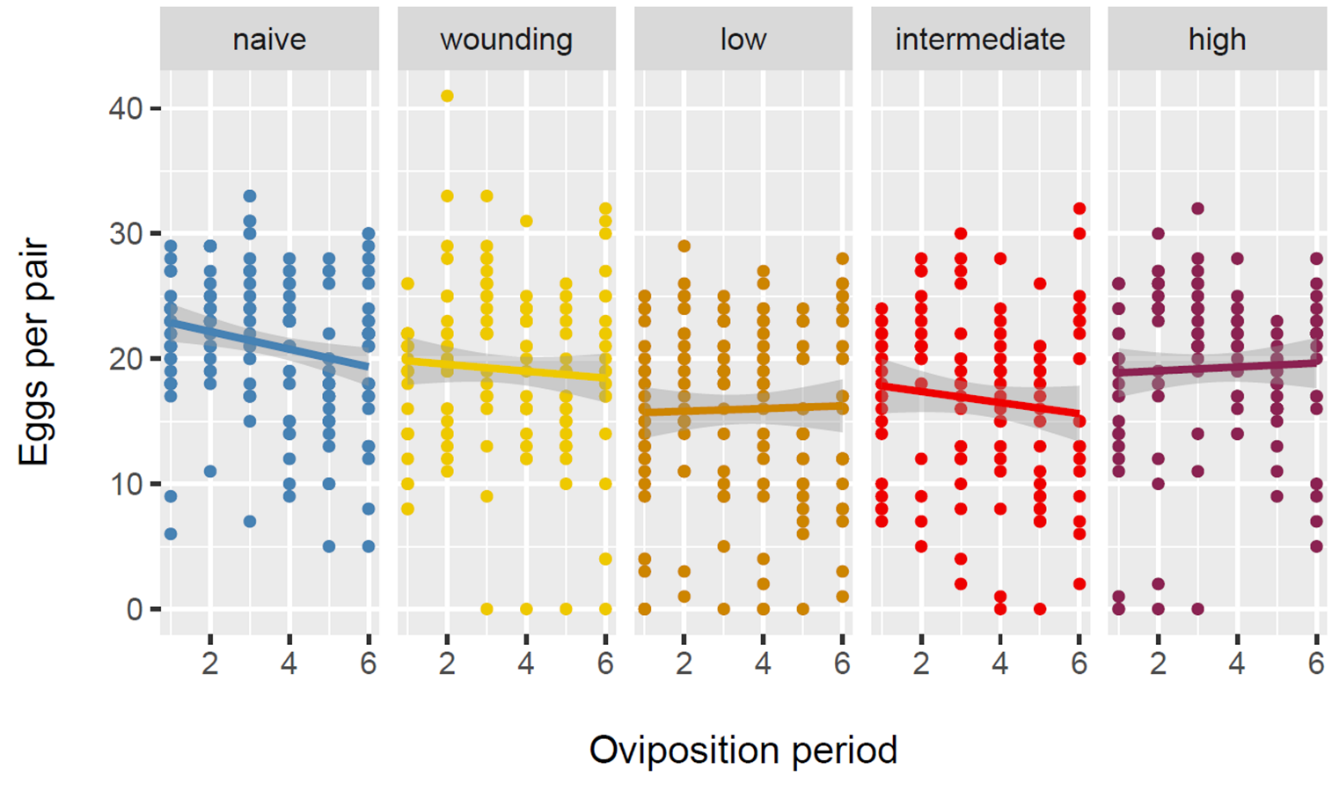

Suppl. Figure 1 Eggs produced in each oviposition period according to maternal exposure treatment (naïve n=25, wounding n=23, low dose n=22, intermediate dose n=20, high dose n=19)

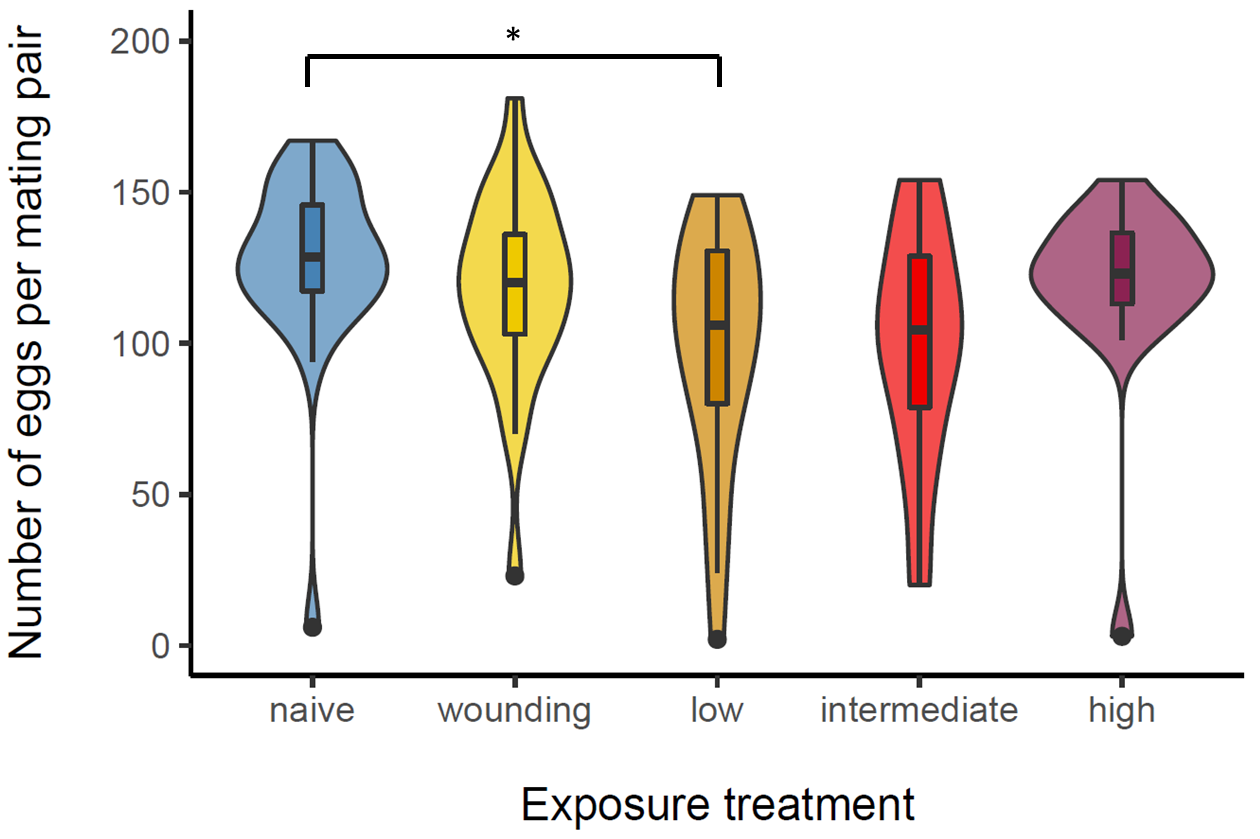

Suppl. Figure 2 Fecundity after maternal exposure (including females that died during the oviposition period. Total number of eggs produced by individual pairs over six 24h oviposition periods (naïve n=26, wounding control n=23, low dose of heat killed Bt n=23, intermediate dose n=20, high dose n=19). Asterisks indicate significant differences (*=p<.05).

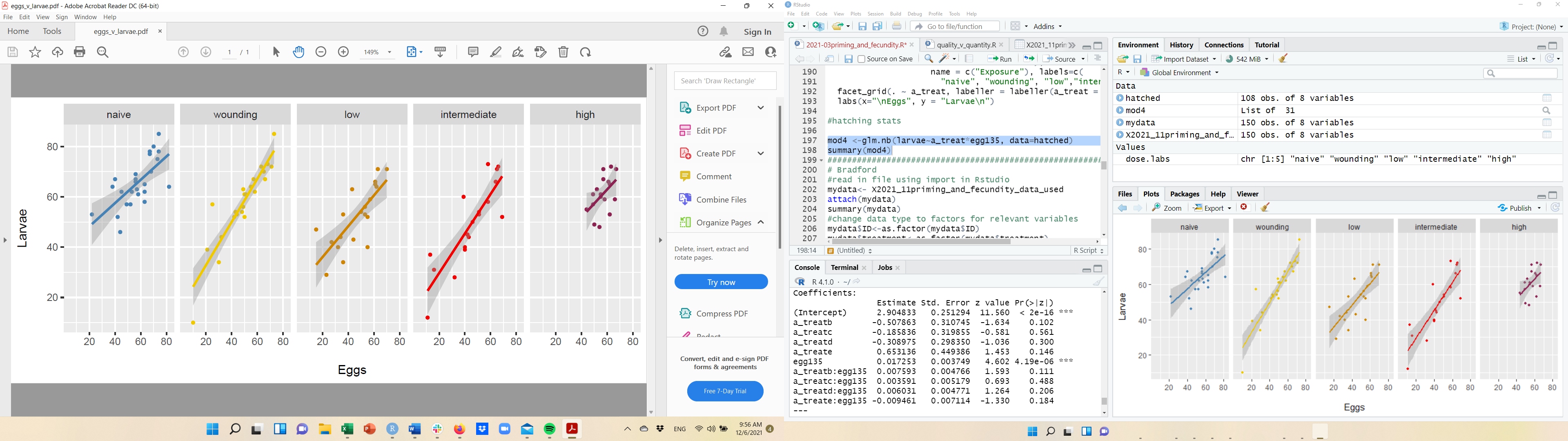

Suppl. Figure 3 Hatching rate. Number of hatched larvae against laid eggs (total numbers forovp 1, 3 and 5) per maternal exposure treatment (naïve n=25, wounding n=23, low dose n=22, intermediate dose n=20, high dose n=18).

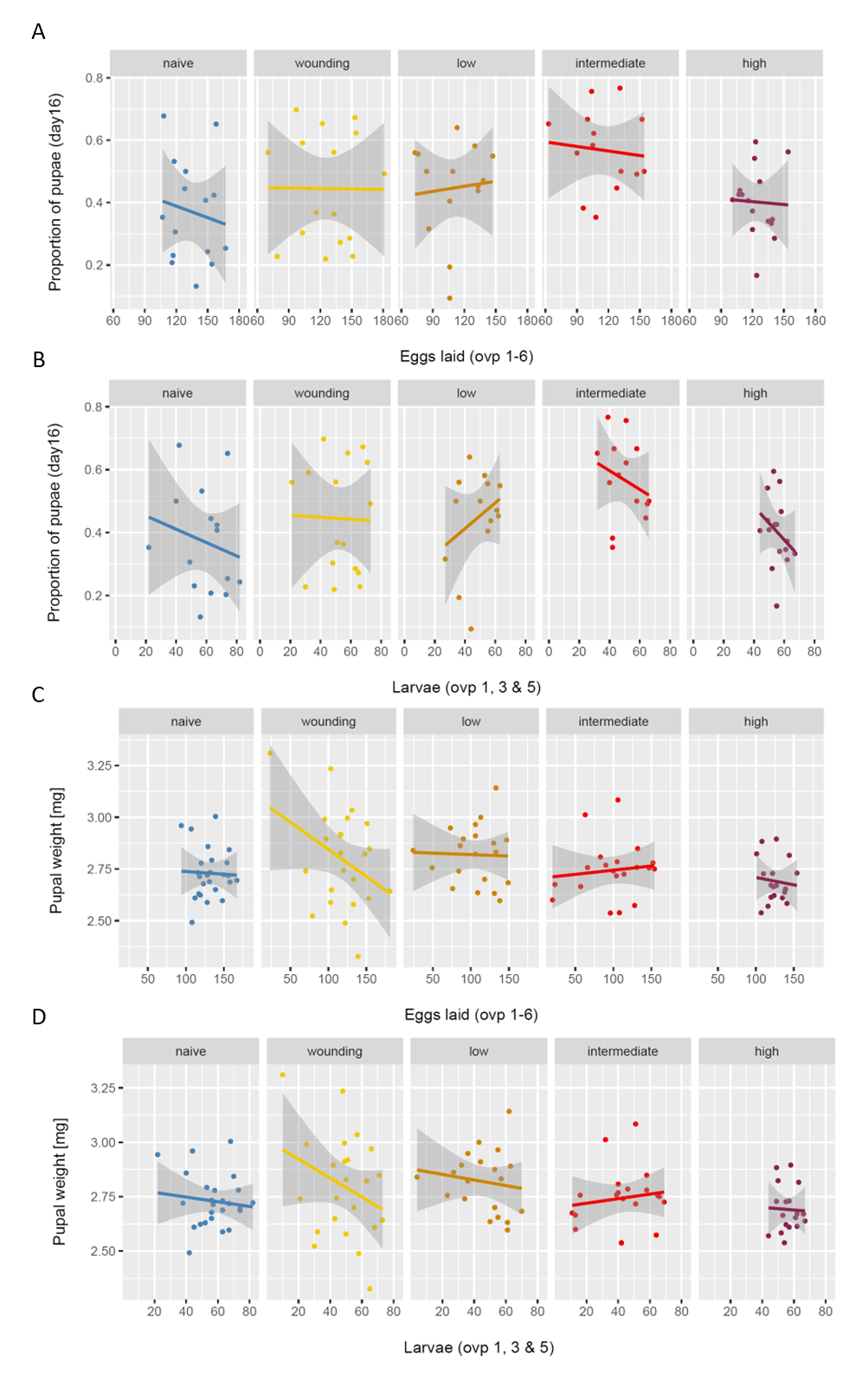

Suppl. Figure 4 Offspring quality-quantity relationship for all five maternal treatment groups (naïve, wounding, low, intermediate and high dose of heat-killed bacteria) A) Total number of laid eggs (all oviposition periods) against number of pupated individuals on day sixteen post oviposition per mating pair B) Total number of hatched larvae (oviposition period 1, 3 and 5) against number of pupated individuals on day sixteen post oviposition per mating pair C) Total number of laid eggs (all oviposition periods) against mean weights calculated for two pupae per mating pair D) Total number of hatched larvae (oviposition period 1, 3 and 5) against mean weights calculated for two pupae per mating pair. None of the correlations were statistically significant (n=18-25).

Suppl. Table 1 Cox Proportional Hazard Analysis for the survival of females after exposure treatment over the six oviposition periods, relative to naïve controls.

| **Treatment**  **(compared to naïve)** | **Hazard ratio** | **SE** | **z-value** | **p-value** |
| --- | --- | --- | --- | --- |
| wounding control | 2.1088 | 1.2248 | 0.609 | 0.5424 |
| low dose | 8.2084 | 1.0607 | 1.985 | 0.0472 |
| intermediate dose | 8.5023 | 1.0692 | 2.002 | 0.0453 |
| high dose | 6.84 | 1.0802 | 1.78 | 0.0751 |
| N= 138, number of events= 24 | | | | |

Suppl. Table 2 Analysis of fecundity measurements (egg counts, larval count, hatching rate). Shown are the results of GLMs or GLMMs for eggcount data from all mothers, egg count data for surviving mothers, egg counts per oviposition period (ovp), number of larvae produced, and hatching rate of larvae from eggs.

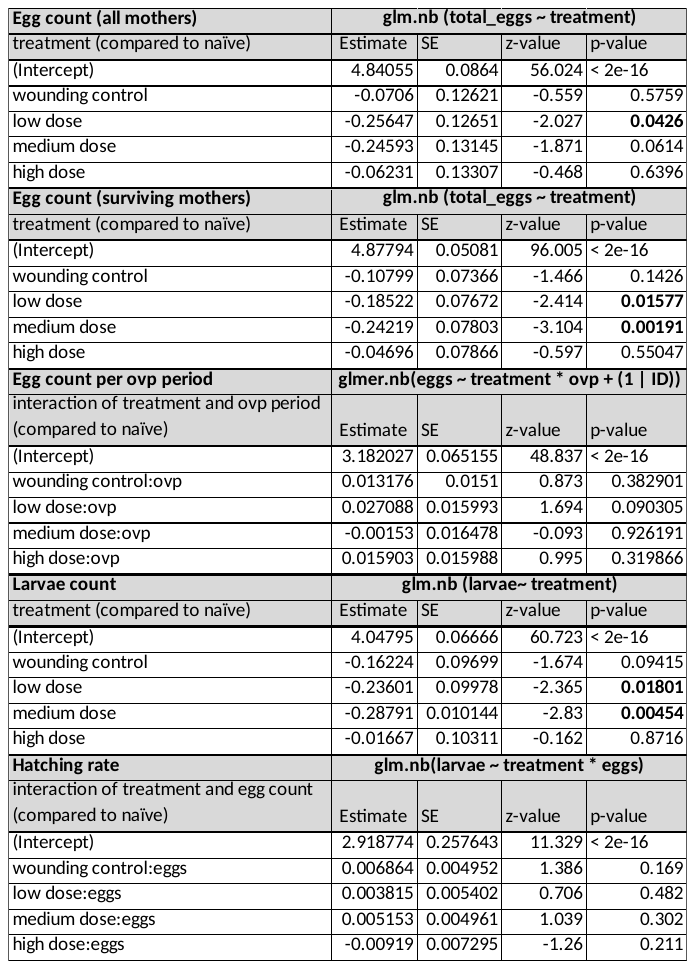

Suppl. Table 3 Analysis of protein content per egg. Shown are the results of a GLMM.

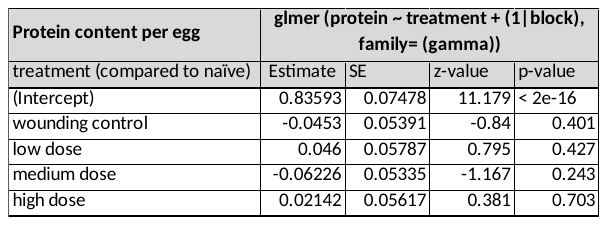

Suppl. Table 4 Analysis of correlations between offspring quantity (produced eggs and larvae) and quality (egg protein content, pupation, and pupal weight)

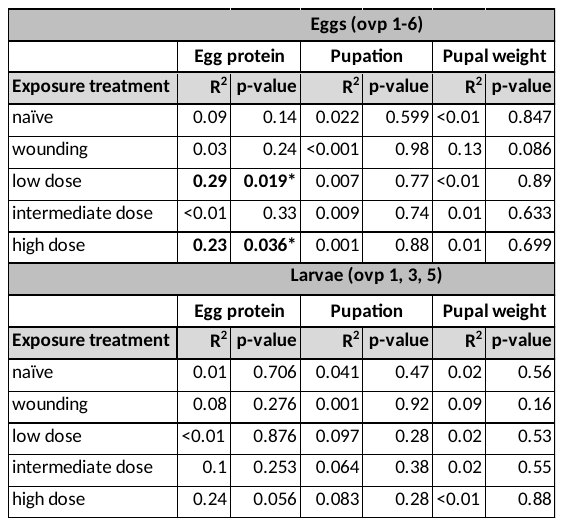

Suppl. Table 5 Analysis of development (proportion of puape on day 16) and pupal weight. Shown are the results of GLMMs.

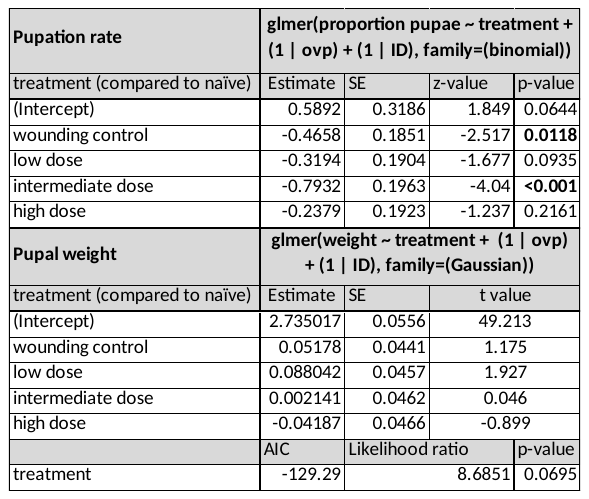

Suppl. Table 6 Percentages of larval offspring that survived a septic Bt infection for at least 48h. Percentages are given per maternal exposure treatment and oviposition period (n=45-68).

| **Exposure treatment** | **Ovp 1** | **Ovp 3** | **Ovp 5** | **mean** |
| --- | --- | --- | --- | --- |
| naive | 82% | 68% | 88% | **79.3%** |
| wounding | 81% | 69% | 86% | **78.6%** |
| low dose | 77% | 66% | 94% | **79%** |
| intermediate dose | 85% | 63% | 98% | **82%** |
| high dose | 92% | 67% | 90% | **83%** |
| **mean** | **83.4%** | **66.7%** | **91.2%** |  |

Suppl. Table 7 Analysis of offspring survival after Bt infection. Shown are the results of a GLM for maternal exposure treatment and oviposition period.

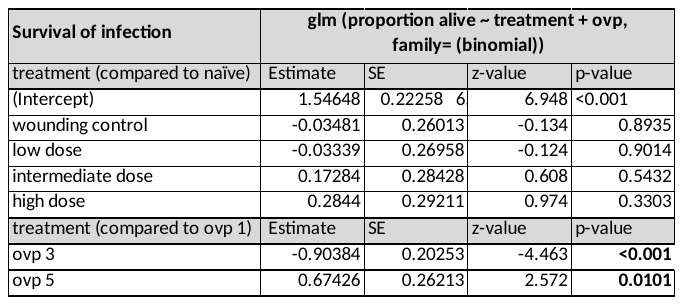
